# Effective Therapy Targeting Cytochrome *bc_1_* Prevents *Babesia* Erythrocytic Development and Protects from Lethal Infection

**DOI:** 10.1101/2021.03.31.438003

**Authors:** Joy E. Chiu, Isaline Renard, Anasuya C. Pal, Pallavi Singh, Pratap Vydyam, Jose Thekkiniath, Madelyn Kumar, Shalev Gihaz, Sovitj Pou, Rolf W. Winter, Rozalia Dodean, Aaron C. Nilsen, Michael K. Riscoe, J. Stone Doggett, Choukri Ben Mamoun

**Affiliations:** Department of Internal Medicine, Section of Infectious Diseases, Yale School of Medicine, New Haven, Connecticut, USA; Veterans Affairs Portland Health Care System, Portland, Oregon, USA

**Author notes:** Joy E. Chiu and Isaline Renard contributed equally to this work. Joy E. Chiu conducted initial *in vitro* and *in vivo* studies. Isaline Renard conducted subsequent *in vivo* efficacy studies using an optimized mouse model.

## Abstract

Targeting conserved metabolic processes that are essential for viability of pathogens, such as *Plasmodium* and *Babesia* that cause blood-borne diseases, is an effective strategy to eliminate malaria and babesiosis infections with no recrudescence. One interesting target is the mitochondrial cytochrome *bc*_*1*_ complex, which could be inhibited by drugs such as endochin-like quinolones (ELQ) and atovaquone. We used the tick-transmitted and culturable blood-borne pathogen *Babesia duncani* to evaluate the structure-activity relationship, safety, efficacy and mode of action of ELQs. We identified a potent and highly selective ELQ prodrug (ELQ-502), which alone or in combination with atovaquone eliminates *B. microti* and *B. duncani* infections *in vitro* and in mouse models of parasitemia and lethal infection. The strong efficacy at low dose, excellent safety, bioavailability and long half-life of this experimental therapy makes it an ideal clinical candidate for the treatment of human infections caused by *Babesia* and its closely related apicomplexan parasites.

## Introduction

Genomic and phylogenetic analyses of several blood-borne pathogens including *Plasmodium* and *Babesia* parasites have offered unique insights into the biology of these closely related organisms and revealed conserved metabolic pathways that could be exploited for the development of multipurpose therapies. The rapid increase in the number of tick-borne diseases reported over the last decade, including human babesiosis cases caused by *Babesia* species, highlights the need for developing new and effective therapies (1, 2). *Babesia microti, Babesia duncani, Babesia divergens* and *Babesia venatorum* are the main species associated with human disease (1-3). Symptoms of severe cases of human babesiosis include acute respiratory distress syndrome, hemolytic anemia, multiple organ failure and possibly death (2, 3). Although transmission of the parasite to human occurs mainly through the bite from an infected tick, increasing number of human-to-human transmission cases through blood transfusion have been reported (4, 5). In 2009, *Babesia* was added to the list of infectious agents identified as potential threats to the safety of blood supplies and it has become a nationally notifiable disease since 2011 (6). Today, *B. microti* is one of the most commonly reported transfusion-transmitted pathogens and is the leading infectious cause of transfusion-related deaths, with one in five cases of babesiosis acquired through blood transfusion resulting in fatal outcome (7, 8).

Due to its shared clinical features with malaria, the current treatments for babesiosis are based on the use of combinations of known antimalarial drugs. These include a combination of atovaquone + azithromycin and a combination of clindamycin + quinine for moderate and severe babesiosis, respectively(1, 9). However, these therapies are often associated with adverse effects or with the development of drug-resistant parasites. As a result, there is a growing need to develop tailored therapies or combination therapies that are specific to *Babesia*, have limited to no side-effects and prevent the development of drug-resistant parasites.

A previous study identified endochin-like quinolones (ELQs) as a novel class of anti-*B. microti* drugs that target the quinone reduction (Q_i_) site of the parasite’s cytochrome *b* (Cytb) (10). Because of the conserved nature of the *Cytb* gene between various apicomplexan parasites, ELQ-based drugs have also shown efficacy against other species such as *Plasmodium falciparum* and *Toxoplasma gondii*, the agents of human malaria and toxoplasmosis, respectively (11-15). However, the poor bioavailability of these compounds due to high crystallinity and low aqueous solubility was a limiting factor to their *in vivo* evaluation, precluding administration of doses higher than 10 mg/kg. To improve the oral bioavailability of this class of compounds, a prodrug strategy was developed whereby the carbonyl of the quinolone core was esterified, resulting in decreased crystallinity of the prodrug and increased plasma concentration of the corresponding drug following cleavage by the host esterase (16). With this improved bioavailability, a new prodrug candidate, ELQ-334, was evaluated *in vivo* against *B. microti*. Although single drug treatment resulted in the emergence of resistant parasites, associated with a mutation in the Cytb Q_i_ site, a combination of ELQ-334 + atovaquone at doses as low as 5 + 5 mg/kg resulted in elimination of *B. microti* infection in 100% of mice (10). However, due to the lack of an *in vitro* culture system for *B. microti*, structure-activity relationship (SAR) analyses to improve the potency and pharmacological properties of ELQ-334 and its parent drug ELQ-316 were limited. The recent breakthrough in the continuous propagation of *B. duncani in vitro* in human red blood cells and the availability of animal models of parasitemia and lethal infection provided a unique opportunity to screen ELQ derivatives for *in vitro* potency and safety and to evaluate their efficacy in mouse models of babesiosis (17).

Here we report successful ELQ SAR studies using this new model system and the identification of ELQ-502 as a highly potent, safe and effective drug, which alone or in combination with atovaquone achieves complete elimination of parasite burden and prevents lethal infection.

## Results

### Structure-activity relationship analysis of ELQ compounds

To identify new ELQ compounds with improved efficacy and selectivity, structure-activity relationship studies were conducted using *B. duncani* propagated *in vitro* in human red blood cells (17). A set of 28 ELQ analogs were synthesized and tested for their ability to inhibit the development of *B. duncani* in human red blood cells (Fig. 1a and b). Of these drugs, 20 compounds inhibited growth of *B. duncani* by more than 80% at 1 µM. Seven of these compounds inhibited growth of the parasite by more than 80% at 100 nM (Fig. 1b). ELQ-447 and its respective prodrug, ELQ-502, were further selected based on their *in vitro* efficacy and desirable safety profile. Both parent drug (ELQ-447) and prodrug (ELQ-502) showed excellent potency *in vitro* (Fig. 1c). With an IC_50_ of 165 ± 1 nM, ELQ-447 displayed a potency similar to that of the previously reported ELQ-316 and ELQ-334, whose IC_50_ values were 136 ± 1 nM and 193 ± 56 nM, respectively (Fig. 1c and d). Interestingly, the *in vitro* potency of ELQ-502 (IC_50_ = 6 ± 2 nM) was 28 x higher than its parent drug (Fig. 1c). We further evaluated the safety profile of ELQ-502 by measuring its toxicity profile in HeLa, HepG2 and HCT116 cell lines under conditions permissive for glycolysis (glucose-based media) or for mitochondrial oxidative phosphorylation (galactose-based media) (Table 1). Under all these conditions, the calculated IC_50_ of ELQ-502 was greater than 5 µM, and its therapeutic index (IC_50_ human cells/IC_50_ *B. duncani*) greater than 833 (Table 1), making it an ideal lead candidate for further evaluation in animal models.

**Figure 1.**
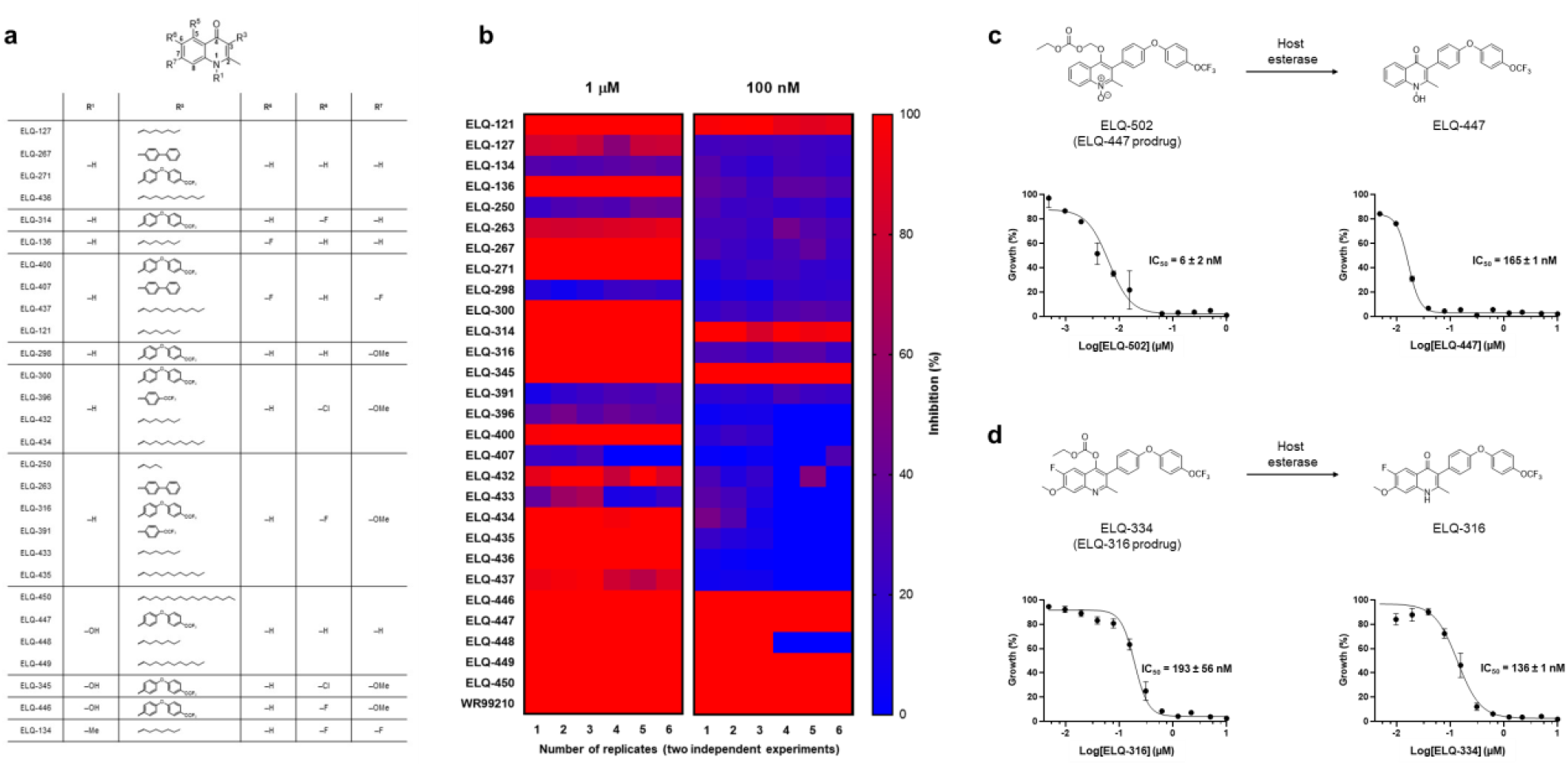
*In vitro* efficacy of ELQ analogs against *B. duncani* in human RBCs. (**a**) Chemical structure of ELQs used in this study. (**b**) Twenty-eight ELQ derivatives were evaluated *in vitro* against *B. duncani* at 1 µM and 100 nM. WR99210, a potent antiparasitic drug(25, 26), was used as a positive control and used to determine 100% growth inhibition. (**c-d**) Structure and potency of ELQ drugs and prodrugs against *B. duncani* and IC_50_ determination: ELQ-447 and ELQ-502 (**c**); ELQ-316 and ELQ-334 (**d**).

**Table 1.** Activity of ELQ-502 in *B. duncani*-infected erythrocytes, HeLa, HepG2 and HCT116 cells.

| Compounds | IC<sub>50</sub><br>*B. duncani* | IC<sub>50</sub><br>HeLa |  |  | IC<sub>50</sub><br>HepG2 |  |  | IC<sub>50</sub><br>HCT116 |  |  | Therapeutic<br>Index |
| --- | --- | --- | --- | --- | --- | --- | --- | --- | --- | --- | --- |
|  |  | H-Glc | L-Glc | Gal | H-Glc | L-Glc | Gal | H-Glc | L-Glc | Gal |  |
| ELQ-502 | 6 ± 2 nM | > 10 µM | > 5 µM | > 10 µM | > 10 µM | > 10 µM | > 10 µM | > 10 µM | > 10 µM | > 5 µM | > 833 |
| Atovaquone | 72 ± 6 nM | > 5 µM | > 5 µM | > 5 µM | > 5 µM | > 5 µM | > 5 µM | > 5 µM | > 5 µM | > 5 µM | > 69 |

### *In vivo* efficacy of ELQ-502 in *B. duncani*-infected mice

The high potency and low toxicity of ELQ-502 against *B. duncani in vitro* led us to investigate the efficacy of this prodrug in mice infected with *B. duncani* WA-1 isolate, both as a single therapy or in combination with atovaquone (Fig. 2). These assays were conducted in C3H/HeJ mice, which show a mortality rate of 100% 10 to 11 days post-infection with 10^3^ *B. duncani*-infected red blood cells. Whereas oral administration of the vehicle alone (PEG-400) resulted in 100% mortality by day 11 post-infection (DPI 11), with parasitemia reaching ∼ 6% (Fig. 2a and e), oral administration of atovaquone, ELQ-502 or ELQ-502 + atovaquone at 10 mg/kg each for 10 days (DPI 1 – 10) resulted in elimination of parasitemia in most if not all infected animals (Fig.2b-d). Complete elimination of *B. duncani* infection with no recrudescence was achieved with ELQ-502 and ELQ-502 + atovaquone (Fig. 2c and d). No animals succumbed to lethal *B. duncani* infection following treatment with ELQ-502 or ELQ-502 + atovaquone (Fig. 2e). Similar to previous analyses of relapse following drug treatment of *B. microti*-infected mice (10), treatment with atovaquone was accompanied by recrudescence in 2 out of 10 mice (Fig. 2b) starting at DPI 20, whereas no parasitemia could be detected in the remaining 8 mice up to DPI 45 (end of study). Surprisingly, analysis of the *Cytb* gene from re-emerging parasites showed lack of mutations in this gene, and *in vitro* culture of parasites that emerged from atovaquone-treated mice showed that they remained susceptible to the drug (Fig. 2f and g).

**Figure 2.**
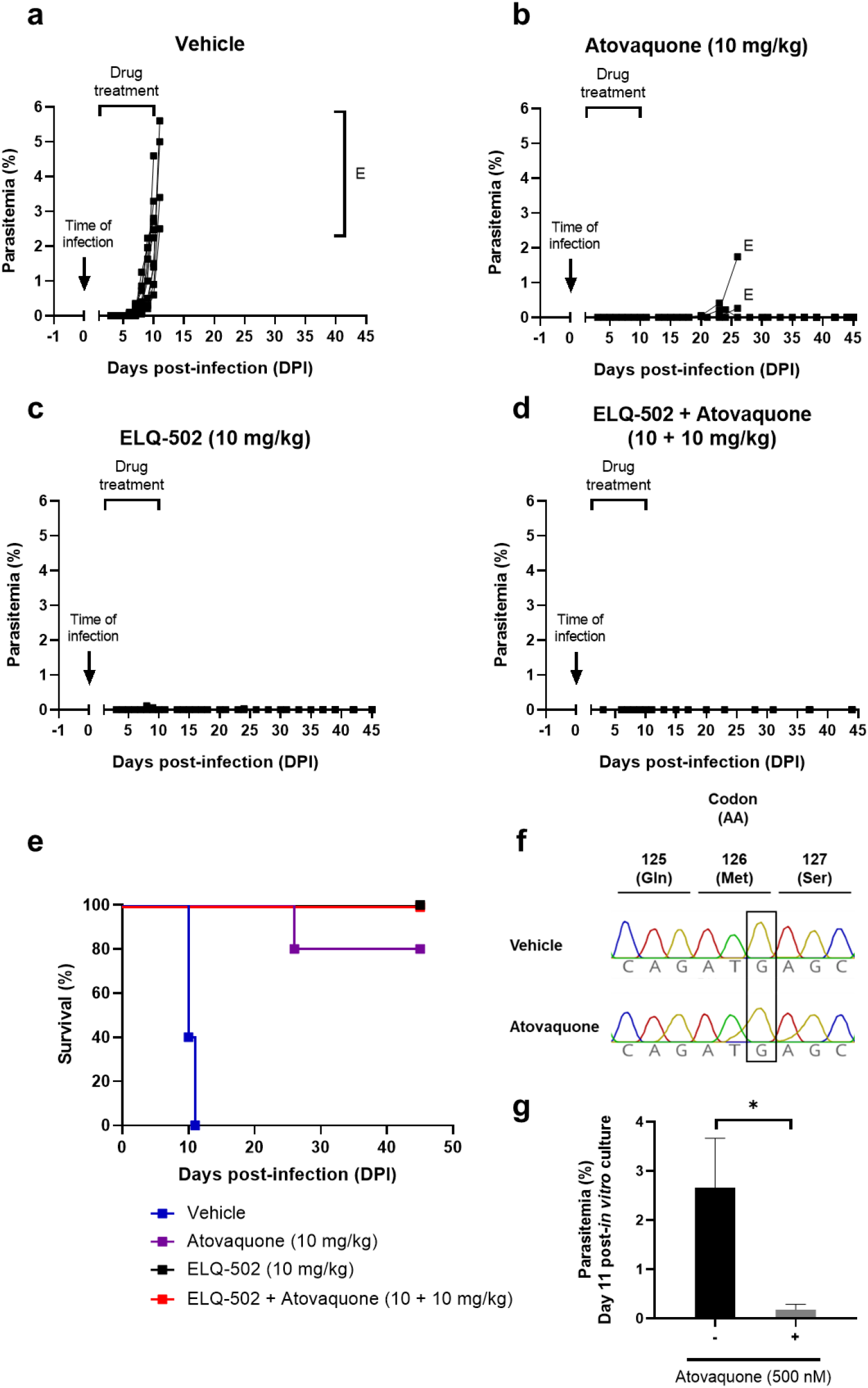
Efficacy of atovaquone and ELQ-502, alone or in combination, in a mouse model of lethal B. duncani infection. (**a-e**) Female C3H/HeJ mice were infected i.v. with 10^3^ *B*.*duncani*-infected red blood cells. Animals were treated daily (DPI 1 – DPI 10) by oral gavage with the vehicle alone (PEG-400) (**a**), atovaquone at 10 mg/kg (**b**), ELQ-502 at 10 mg/kg (**c**) or a combination of ELQ-502 + atovaquone at 10 + 10 mg/kg (**d**). E: indicates when an individual mouse was euthanized. (**e**) Survival rate of *B. duncani*-infected mice in the absence or following treatment with atovaquone, ELQ-502 or atovaquone + ELQ-502. (**f**) Representative sequencing chromatogram of recrudescent parasites from atovaquone-treated *B. duncani*-infected mice. Genomic DNA was extracted from recrudescent parasites and used to amplify the *BdCytb* gene followed by sequencing. As a control, the chromatogram of wild type parasites (obtained from control animals) is shown. No mutation in the *BdCytb* gene could be found. (**g**) *In vitro* drug susceptibility of recrudescent parasites from atovaquone-treated *B. duncani*-infected mice. Parasites were cultured *in vitro* in human erythrocytes in the absence or presence of atovaquone at 500 nM. At day 11 post-*in vitro* culture, parasites cultured in the absence of drug reached ∼2.5% parasitemia, whereas parasites treated with atovaquone remained at 0.18% parasitemia. *P < 0.02.

### *In vivo* efficacy of ELQ-502 in *B. microti*-infected mice

Following the promising results observed in the *in vivo* model of *B. duncani*, ELQ-502 was subsequently evaluated alone or in combination with atovaquone in *B. microti*-infected mice using the SCID mouse model of *microti* infection. Whereas parasitemia in vehicle-administered control animals reached high levels by DPI 20 and leveled out at 60-80% parasitemia for the remainder of the study (Fig. 3a), treatment with atovaquone at 10 mg/kg was accompanied by the emergence of recrudescent parasites by DPI 24 (Fig. 3b). Interestingly, *B. microti*-infected mice treated with ELQ-502 alone (10 mg/kg) or in combination with atovaquone (10 + 10 mg/kg) showed no sign of infection for at least 35 days after completion of the drug treatment (Fig. 3c and d) (mice from the combination treatment were monitored for up to 120 days (110 post-treatment) and no parasites could be detected).

**Figure 3.**
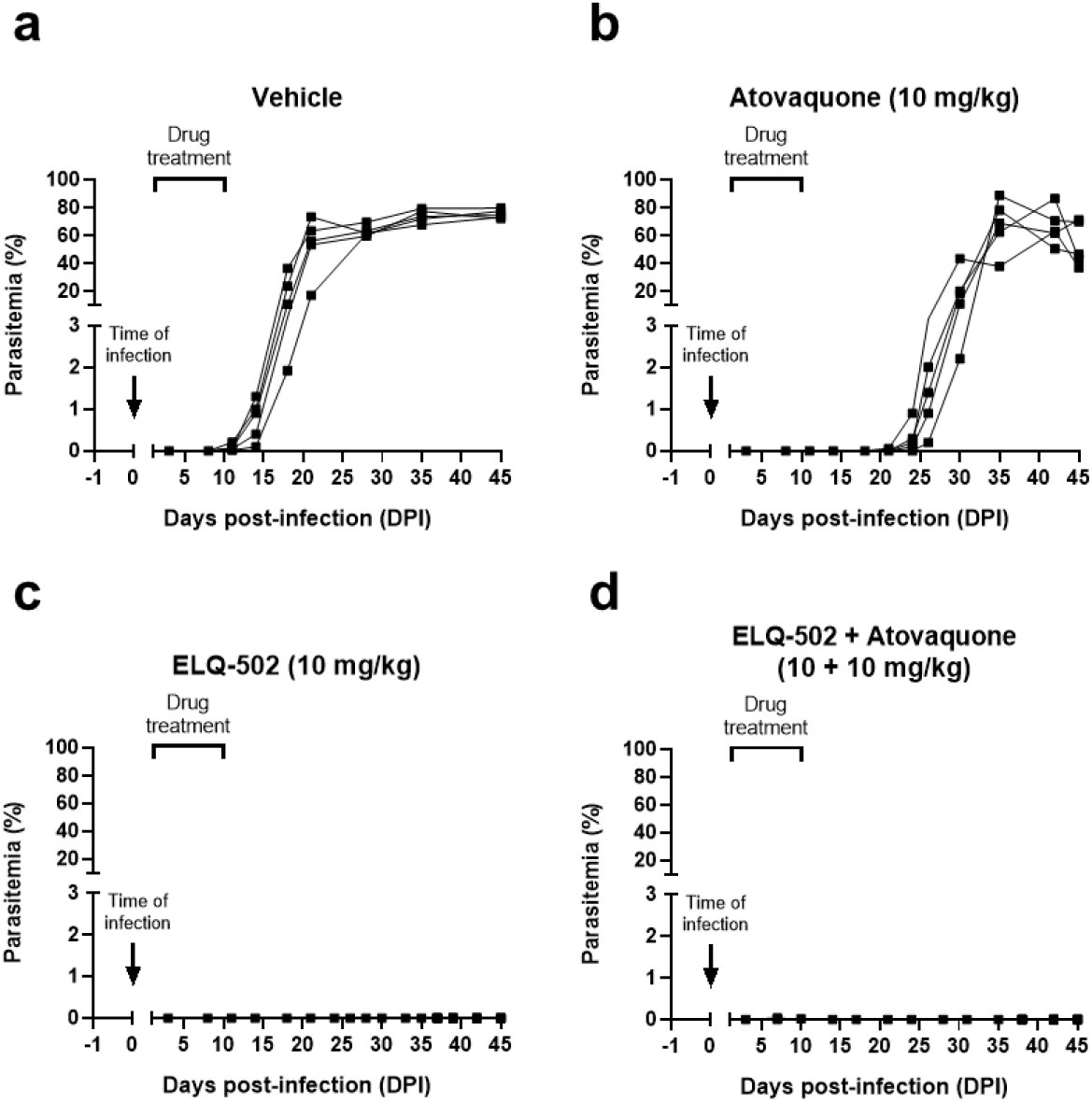
*In vivo* evaluation of atovaquone and ELQ-502, as single drugs or in combination in *B. microti*-infected mice. Female SCID mice were injected with either 10^4^ i.v. or 10^6^ i.p. *B*.*microti*-infected red blood cells. Animals were treated daily (DPI 1 – DPI 10) by oral gavage with the vehicle alone (PEG-400) (**a**), atovaquone at 10 mg/kg (**b**), ELQ-502 at 10 mg/kg (**c**) or ELQ-502 + atovaquone at 10 + 10 mg/kg (**d**).

### Evidence of a dual mode of action of ELQ-502 through drug-drug interaction analyses

The ability of ELQ-502 to achieve cure with no recrudescence in both *B. duncani*- and *B. microti*-infected mice following a 10-day treatment at 10 mg/kg starting at D1 post-infection led us to further investigate its mode of action. This was achieved by examining the possible additive, synergistic or antagonistic effect of dual combinations of ELQ-502 with drugs known to inhibit either the Q_o_ site (atovaquone) or the Q_i_ site (ELQ-331 and ELQ-468) of the cytochrome *bc*_*1*_ complex (18). Both ELQ-331 and ELQ-468 were found to have potent activity against *B. duncani in vitro* with IC_50_ values of 141 ± 22 nM and 15 ± 1 nM, respectively (Fig. 4a and b). Interestingly, combinations of ELQ-502 with either atovaquone, ELQ-331 or ELQ-468 were found to be synergistic with a mean fractional inhibitory concentration (ΣFIC_50_) value of 0.6, 0.4 and 0.9, respectively (Fig. 4c-e and h). As a control, a combination of ELQ-334 and atovaquone (Fig. 4f and h) was also found to be synergistic, consistent with its previously reported efficacy in *B. microti*-infected mice (10). However, unlike the synergistic effects seen with ELQ-502, a combination of ELQ-331 and ELQ-468 was mainly additive (ΣFIC_50_ = 1.2) (Fig. 4g and h). Together these data suggest that ELQ-502 may target not the only the Q_i_ site of the cytochrome *bc*_*1*_ complex, but also its Q_o_ site and this dual activity may account for its high potency (Fig. 4i).

**Figure 4.**
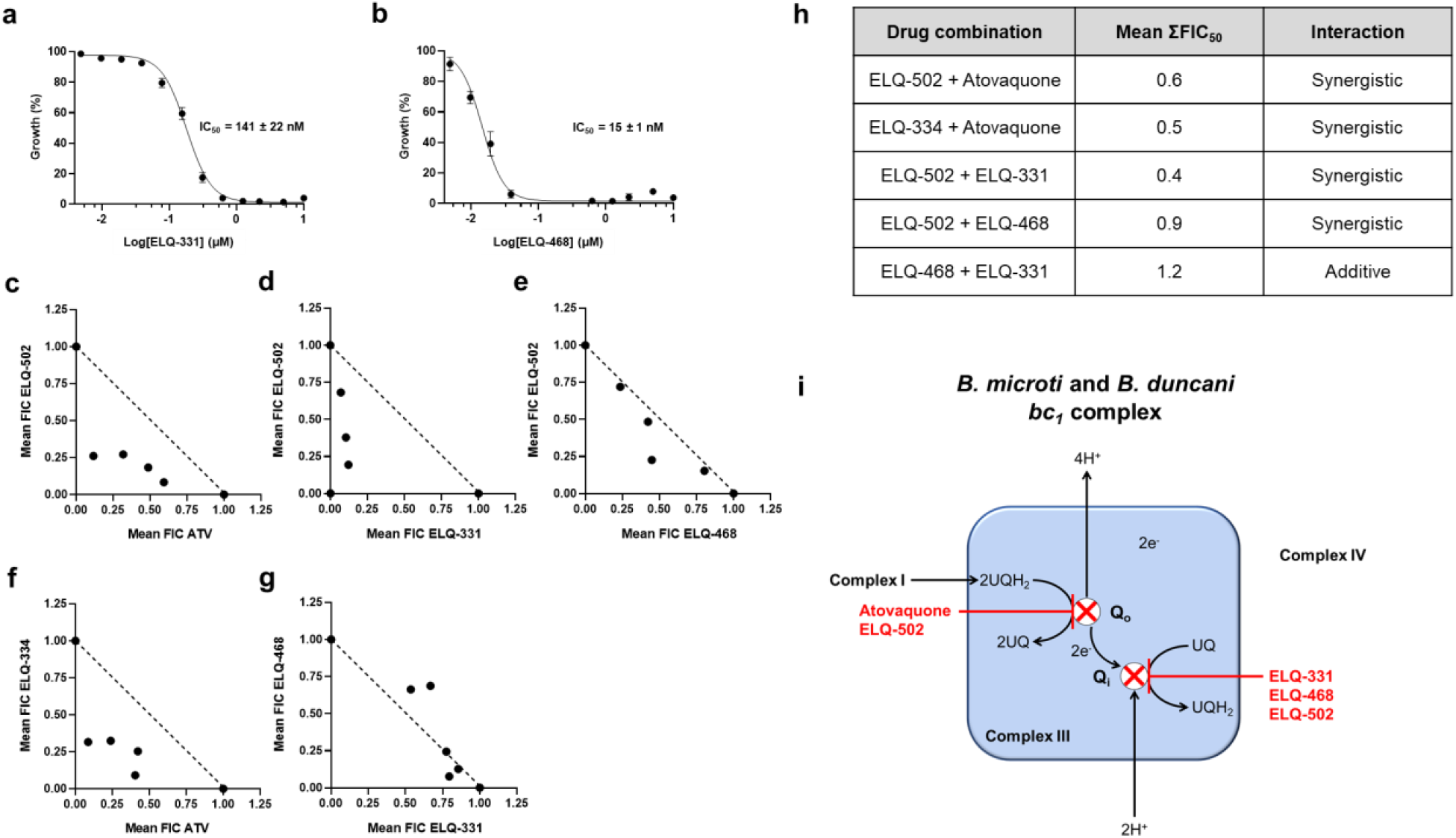
Atovaquone-ELQ and ELQ-ELQ drug-drug interactions. (**a-b**) Potency of two other promising ELQ prodrugs against *B. duncani* and IC_50_ determination: ELQ-331 (**a**) and ELQ-468 (**b**). (**c-g**) Isobolograms depicting the activity of ELQ-502 in combination with atovaquone (**c**), ELQ-331 (**d**) and ELQ-468 (**e**). For reference, isobolograms of combination of ELQ-334 + atovaquone (**f**) and ELQ-331 + ELQ-468 (**g**) are shown. Each point in the graph represents the mean FIC_50_ value from a fixed drug ratio as described in Materials and Methods. A dotted line plotted between the individual mean FIC_50_ value of each drug serves as additive line. (**h**) Summary of drug interactions. (**i**) Schematic representation of the *B. microti* and *B. duncani bc*_1_ complex and proposed mode of action of atovaquone, ELQ-331, ELQ-468 and ELQ-502. IMS: inter-membrane space, MM: mitochondrial matrix.

## Discussion

Our results show that ELQ-502 is a potent and highly effective endochin-like quinolone against *Babesia microti* and *Babesia duncani in vitro* and in mouse models of high parasitemia and lethal babesiosis infection, either as a monotherapy or in combination with atovaquone. ELQs have been shown to target the Q_i_ site or the Q_o_ site of the cytochrome *b* of multiple apicomplexan parasites such as *Plasmodium spp*. (15, 16, 19-21), *Toxoplasma gondii* (11) and *Babesia microti* (10). Whereas the ability to maintain *P. falciparum* and *T. gondii* in culture made it possible to conduct structure-activity relationship studies to identify inhibitors suitable for each pathogen, detailed structure-activity relationship evaluation could not be conducted in *B. microti* due to the lack of an *in vitro* culture system. The recent development of a continuous *in vitro* culture system for *B. duncani* in human red blood cells (17) provides a means to probe different moieties that define the selective potency of ELQs both *in vitro* and in a small animal model.

With an IC_50_ value of 6 ± 2 nM, ELQ-502 was found to be more active than its parent compound, ELQ-447 (IC_50_ = 165 ± 1 nM), and more active than the reference compounds ELQ-316 and ELQ-334 (IC_50_ = 136 ± 1 nM and IC_50_ = 193 ± 56 nM, respectively). This enhanced efficacy can most likely be attributed to the higher aqueous solubility of ELQ-502. Moreover, the lack of activity of ELQ-502 in HepG2, HeLa and HCT116 human cell lines at concentrations ≤ 5 µM resulted in a highly desirable therapeutic index (> 833).

Unlike the previously examined antibabesial prodrug ELQ-334, the prodrug moiety selected to synthesize ELQ-502 was a methoxy ethyl carbonate. The two derivatives share a position 3 diarylether side-chain substituted in *para* of the external ring with a trifluoromethoxy functional group. The use of this type of side-chain was previously reported to improve potency and metabolic stability when assessed against *Plasmodium* parasites (13). One notable difference between ELQ-502 and ELQ-334 is the substitution of the amine with a hydroxyl functional group in the case of ELQ-502, potentially resulting in increased hydrogen bonding and/or electrostatic interactions.

Our studies in both *B. duncani* and *B. microti*-infected mice showed strong *in vivo* efficacy of ELQ-502 as a monotherapy or in combination with atovaquone, resulting in complete parasite elimination with no recrudescence, and in the case of *B. duncani*, 100% survival of animals from lethal infection. To determine whether the success of the ELQ-502 + atovaquone combination therapy *in vivo* can be attributed to synergy between the two drugs, combination studies were carried out *in vitro* in *B. duncani* and identified synergistic interactions between ELQ-502 and atovaquone. To further understand the mode of action of ELQ-502, drug-drug interaction studies were carried out with two ELQ derivatives, ELQ-331 and ELQ-468 with potent activity against *B. duncani in vitro* (IC_50_ = 141 ± 22 nM and IC_50_ = 15 ± 1 nM, respectively). These studies showed synergy between ELQ-502 and either ELQ-331 or ELQ-468, whereas only additive interactions were observed between ELQ-331 and ELQ-468. One possible explanation for the unique potency of ELQ-502 could be that this drug targets both the Q_i_ and Q_o_ sites of the cytochrome *bc*_*1*_ complex. This ability of some ELQ compounds to target both sites has previously been demonstrated for ELQ-400 in yeast (22). Nevertheless, a 5-day treatment with ELQ-502 starting at day 3 resulted in emergence of resistant parasites carrying a mutation in the Q_i_ site. This emphasizes the importance of a combination therapy as an ideal strategy for the development of new treatment for human babesiosis.

In conclusion, we propose that a combination of ELQ-502 + atovaquone might offer a superior option to current treatments for human babesiosis, which are associated with adverse events, low efficacy and resistance occurrence. Future studies aimed to characterize the safety of this compound in other mammals and humans are needed before it could be evaluated for efficacy in human trials.

## Materials and Methods

### Chemistry

Unless otherwise stated, chemicals were purchased from commercial suppliers and used as received. ELQ compounds were synthesized following methods previously described by Doggett *et al*. (11), Nilsen *et al*. (13) and Frueh *et al*. (18), identified by proton nuclear magnetic resonance (^1^H NMR), and determined to be ≥ 95% pure by reversed-phase high-performance liquid chromatography (RP-HPLC).

### *In vitro* culture of *B. duncani*

*In vitro* culture of *B. duncani* was carried out as previously reported by Abraham *et al*. (17). A cryostock of *B. duncani* was thawed at 37°C. The content of the vial (≈ 1 mL) was transferred to a 50 mL falcon tube and 200 µL of 12% NaCl (w/v) were added dropwise. The resulting mixture was incubated at room temperature for 5 min. 10 mL of 1.6% NaCl (w/v) were slowly added and the suspension was centrifuged at 1800 rpm for 5 min. The supernatant was removed leaving ≈ 500 µL to resuspend the pellet. 10 mL of pre-warmed complete HL-1 medium (HL-1 base medium (Lonza, 344017) supplemented with 20% heat-inactivated FBS (Sigma, F4135), 2% 50X HT Media Supplement Hybri-Max™ (Sigma, H0137), 1% 200 mM L-Glutamine (Gibco, 25030-081), 1% 100X Penicillin/Streptomycin (Gibco, 15240-062) and 1% 10 mg/mL Gentamicin (Gibco, 15710-072)) were added slowly and the resulting mixture was centrifuged at 1800 rpm for 5 min. The supernatant was removed, and the pellet was resuspended in 500 µL of complete HL-1 medium. The parasite suspension and 100 µL of 50% hematocrit A^+^ RBCs (5% final hematocrit) were plated in one well of a 24-well plate. The final volume was made up to 1 mL by adding complete medium. The plate was incubated at 37°C under a 2% O_2_ / 5% CO_2_ / 93% N_2_ atmosphere in a humidified chamber. Culture medium was changed daily and parasitemia was monitored by light microscope examination of Giemsa-stained thin-blood smears.

### *In vitro* drug efficacy in *B. duncani*-infected human red blood cells

Experiments were performed to evaluate the effect of ELQs on intra-erythrocytic development cycle (IDC) inhibition of *B. duncani* and determine their IC_50_ values. *In vitro* parasite culture (0.1% parasitemia with 5% hematocrit in HL-1 medium) was treated with decreasing concentrations (two-fold dilution starting from 10 µM) of the compound of interest for 60 h in a 96-well plate. After treatment, parasite viability was determined by SYBR Green-I method(17). Briefly, to 100 µL of parasite culture, an equal volume of lysis buffer (0.008% saponin, 0.08% Triton-X-100, 20 mM Tris-HCl (pH = 7.5) and 5 mM EDTA) containing SYBR Green-I (0.01%) was added and incubated at 37°C for 1 h in the dark. The fluorescent intensity was measured for λ_ex_ = 480 nm and λ_em_ = 540 nm, using a BioTek Synergy™ Mx Microplate Reader. Background fluorescence (uninfected RBCs in HL-1 medium) was subtracted from each measurement. The half-maximal inhibitory concentration (IC_50_) was determined from a sigmoidal dose-response curve using GraphPad Prism version 8.1. Data are shown as mean ± SD of two independent experiments performed in biological triplicates.

### Drug activity in mammalian cells

HeLa, HepG2 and HCT116 cells were purchased from ATCC. Cell toxicity studies were performed at the Yale Center for Molecular Discovery using seeding cell densities optimized by the Center: HeLa (400 cells/well), HepG2 (1500 cells/well) and HCT116 (1500 cells/well). Each cell line was grown in three different media: high-glucose medium (high-glucose Dulbecco’s modified Eagle’s medium (DMEM) (Invitrogen 11995-065) containing 25 mM glucose, 1 mM sodium pyruvate and supplemented with 5 mM HEPES, 10% FBS and penicillin-streptomycin (50 U/mL penicillin, 50 µg/mL streptomycin), low-glucose medium (DMEM deprived of glucose (Invitrogen 11966-025) supplemented with 5.5 mM glucose, 5 mM HEPES, 10% FBS, 1 mM sodium pyruvate and penicillin-streptomycin as above) and galactose medium (DMEM deprived of glucose (Invitrogen 11966-025) supplemented with 10 mM galactose, 2 mM glutamine, 5 mM HEPES, 10% FBS, 1 mM sodium pyruvate and penicillin-streptomycin as above). Cells were plated into 384-well tissue-culture treated plates (Corning 3764) and allowed to adhere overnight prior to compound treatment. On day 2, compounds and DMSO were added using Echo acoustic dispenser (Labcyte), and 72 h later viable cells were detected using CellTiter Glo (Promega). Compounds were tested in a 16-point dose response (10 µM highest final concentration and 2-fold dilutions). Each plate included 16 wells treated with 10% DMSO (positive control for complete toxicity) and 16 wells treated 0.1% DMSO (negative (vehicle) control). Signal-to-background (S/B), coefficient of variation (CV) and Z prime factor (Z’) were calculated for each cell line using mean and standard deviation values of the positive and negative controls to ensure assay robustness. Compound data was normalized to the mean of 10% DMSO wells (set as 100% toxicity) and mean of the vehicle control wells (set as 0% toxicity). Dose-response curves were plotted using GraphPad Prism version 8.1.

### Mouse strains

SCID mice (C.B-17/IcrHsd-Prkdc^scid^) were obtained from Envigo. C3H/HeJ mice were obtained from The Jackson Laboratory. All animal experiments followed Yale University institutional guidelines for care and use of laboratory animals and were approved by the Institutional Animal Care and Use Committees (IACUC) at Yale University.

### *In vivo* drug efficacy in *B. duncani* and *B. microti*-infected mice

Female C3H/HeJ mice (5-6 weeks old, n = 5-10 mice / group) were infected i.v. with 10^3^ *B. duncani* (WA1)-infected RBCs. Treatment was administered daily for 10 days by oral gavage, beginning on day 1 post-infection (DPI 1). Animals received 100 µL of the vehicle alone (PEG-400), atovaquone (10 mg/kg), ELQ-502 (10 mg/kg) or a combination of ELQ-502 + atovaquone (10 + 10 mg/kg). Parasitemia was monitored by Giemsa staining of thin-blood smears.

Female SCID mice (5-6 weeks old, n = 5 mice / group) were infected with either 10^4^ i.v or 10^6^ i.p. *B. microti* (LabS1)-infected RBCs. Treatment was administered daily for 10 days by oral gavage, beginning on day 1 post-infection (DPI 1). Animals received 100 µL of the vehicle alone (PEG-400), atovaquone (10 mg/kg), ELQ-502 (10 mg/kg) or a combination of ELQ-502 + atovaquone (10 + 10 mg/kg). Parasitemia was monitored by Giemsa staining of thin-blood smears.

### Genomic DNA isolation, *Cytb* PCR and sequencing

Blood was collected and allowed to clot at room temperature for 30 min. Genomic DNA was extracted from the blood pellet using DNeasy Blood and Tissue kit (QIAGEN, 69506). PCR was performed on the extracted DNA samples to amplify the *B. duncani* cytochrome b gene. The following primers were used: 1F, 5’–GGATACAGGGCTATAACCAACAA –3’ and 2R, 5’– GGAAGTTGGCGTCTAGAGTCACTC–3’ and thermocycling conditions were as follows: 95°C for 5 min, 12 x [95°C for 30 s, 65°C for 30 s, 72°C for 30 s], 33 x [95°C for 30 s, 56°C for 30 s, 72°C for 30 s]. The PCR products were purified using QIAquick PCR Purification Kit (QIAGEN, 28106) and sent for Sanger sequencing to the Keck Sequencing Facility (Yale University) using the previous primers in addition to 1R, 5’ – TATGCAATTTGAAGTGAAATTCC – 3’ and 2F, 5’ – TTGGGTTGGAGACTTTGTCAG – 3’. Sequences were analyzed using Geneious 11.1.5 (https://www.geneious.com).

### *In vitro* evaluation of drug combinations in *B. duncani*-infected human red blood cells

To understand the type of interaction between the drugs of interest in this study, we followed an adapted protocol of a fixed ratio drug combination experiment, a well-established assay in the field of *Plasmodium* (23). Combinations of the drugs of interest were performed using six fixed ratios (5:0, 4:1, 3:2, 2:3, 1:4 and 0:5). For each single drug (5:0 and 0:5), the starting concentration was fixed at 8x IC_50_. For each ratio, a two-fold dilution series was carried out so that the IC_50_ of each individual drug falls in either the third or fourth dilution. The experiment was set up in a 96-well plate, where each well contained 200 µL of complete HL-1 medium with or without drug and 0.1% parasitemia with 5% hematocrit. Plates were incubated at 37°C for 60 h. Parasite viability was determined by SYBR Green-I method as described above. For each ratio, the half-maximal inhibitory concentration (IC_50_) was determined from a sigmoidal dose-response curve. The fractional inhibitory concentration (FIC_50_) of each drug at different ratios was calculated using the following equation:

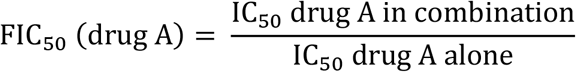

The interaction between drug A and drug B was determined by the sum of FIC_50_ (ΦFIC_50_) using the following formula:

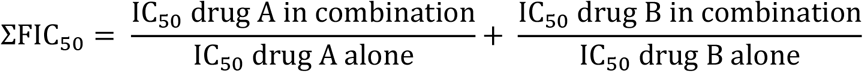

According to the literature, ΣFIC_50_ < 0.8 represents synergy between the two drugs, 0.8 ≤ ΣFIC_50_ < 1.4 suggests additive effect and mean ΣFIC_50_ ≥ 1.4 represents antagonistic properties(24). To illustrate these interactions, isobolograms were plotted as FIC(drug A) = f(FIC(drug B)). Data are averaged from at least two independent experiments, each run in triplicates. Analysis was carried out using GraphPad Prism version 8.1.

## Statistical analysis

Datasets were analyzed with GraphPad Prism version 8.1. Statistical significance was determined using unpaired t-test.

## Acknowledgements

The research described herein was supported by NIH grant R01AI123321 to CBM, JSD and MKR. CBM research is also supported by grants R01AI138139, R01AI152220 and R01AI136118 from NIH and the Steven and Alexandra Cohen Foundation (Lyme 62 2020). Portions of this work were also supported by funds from the United States Department of Veterans Affairs, Veterans Health Administration, Office of Research and Development Program Award number i01 BX003312 (MKR). Additional funding was provided by NIH under award numbers R01AI100569 and R01AI141412 (MKR) and by the U.S. Department of Defense Peer Reviewed Medical Research Program (Log # PR181134) (MKR). The authors would like to thank Yulia Surovtseva and Sheila Umlauf for their help in performing the toxicity studies.

